# Shared input and recurrency in neural networks for metabolically efficient information transmission

**DOI:** 10.1101/2023.03.13.532471

**Authors:** Tomas Barta, Lubomir Kostal

## Abstract

Shared input to a population of neurons induces noise correlations, which decreases the information carried by a population activity. Inhibitory feedback in recurrent neural networks can reduce the noise correlations and thus increase the information carried by the averaged population activity. However, the activity of inhibitory neurons is costly. This inhibitory feedback decreases the gain of the population. Thus, depolarization of its neurons requires stronger excitatory synaptic input, which is associated with higher ATP consumption. Given that the goal of neural populations is to transmit as much information as possible at minimal metabolic costs, it is unclear whether the increased information transmission reliability provided by inhibitory feedback compensates for the additional costs. We analyze this problem in a network of leaky integrate-and-fire neurons receiving correlated input. By maximizing mutual information with metabolic cost constraints, we show that there is an optimal strength of recurrent connections in the network, which maximizes the value of mutual information-per-cost. For higher values of input correlation, the mutual information-per-cost is higher for recurrent networks with inhibitory feedback compared to feedforward networks without any inhibitory neurons. Our results, therefore, show that the optimal synaptic strength of a recurrent network can be inferred from metabolically efficient coding arguments and that decorrelation of the input by inhibitory feedback compensates for the associated increased metabolic costs.

## 1. Introduction

The efficient coding hypothesis poses that neurons evolved due to evolutionary pressure to transmit information as efficiently as possible [1]. Moreover, the brain has only a limited energy budget, and neural activity is costly [2, 3]. The metabolic expense associated with neural activity should, therefore, be considered, and neural systems likely work in an information-metabolically efficient manner, balancing the trade-off between transmitted information and the cost of the neural activity [4, 5, 6, 7, 8].

The principles of information-metabolically efficient coding have been successfully applied to study the importance of the excitation-inhibition balance in neural systems. It has been shown that the mutual information between input and output per unit of cost for a single neuron is higher if the excitatory and inhibitory synaptic currents to the neuron are approximately equal if the source of noise lies in the stochastic nature of the voltage-gated Na^+^and K^+^channels [9]. In a rate coding scheme, where the source of noise lies in the random arrival of pre-synaptic action potentials, the mutual information per unit of cost has been shown to be rather unaffected by the increase of pre-synaptic inhibition associated with an excitatory input [10].

However, the balance of excitation and inhibition is likely to be more important in the context of recurrent neural networks than in the context of single neurons. In recurrent neural networks, the inhibitory input to neurons associated with a stimulus [11] arises as inhibitory feedback from a population of inhibitory neurons. The inhibitory feedback prevents a self-induced synchronization of the neural activity [12] and reduces noise correlations induced by shared input to neurons in the population [13, 14, 15]. Noise correlations are detrimental to information transmission by neural populations [16, 17] and information is likely transmitted by the activity of a population of neurons instead of a single neuron [18]. Therefore, when studying the effect of excitation-inhibition balance on information transmission, it is essential to consider the context of neural populations.

Several studies have analyzed the effect of noise correlations on information transmission properties [16, 17, 19]. However, these studies did not analyze the relationship between the noise correlations and the metabolic cost of neural activity. In our work, we consider a computational model of a small part of the sensory cortex and the noise correlations caused by shared connections from an external thalamic population. The noise correlations may then be reduced by inhibitory feedback, which, however, increases the cost of the neural activity [10]. Our point of interest is the trade-off between improved information transmission due to lower noise correlations and the increase in metabolic costs due to stronger inhibitory feedback.

## 2. Methods

### 2.1. Network model

We modeled a network consisting of three subpopulations: external (ext), excitatory (exc), and inhibitory (inh). The external subpopulation consisted of Poisson neurons, defined by their firing intensity 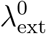 (same for all the neurons in the subpopulation). Neurons in the excitatory and inhibitory subpopulations were modeled as leaky integrate-and-fire (LIF) neurons:

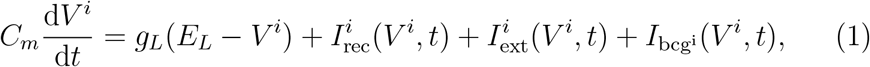

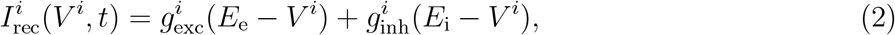

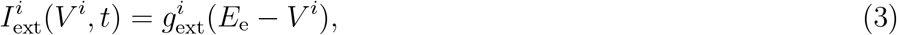

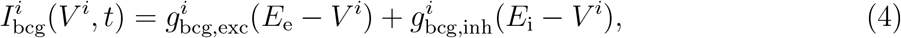

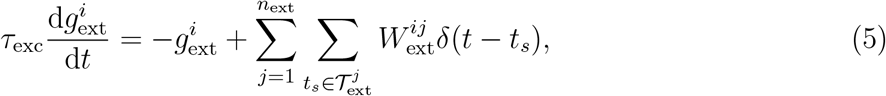

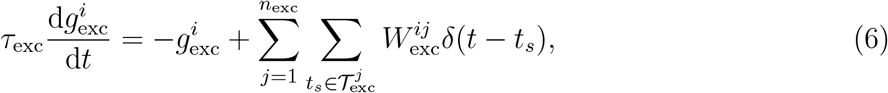

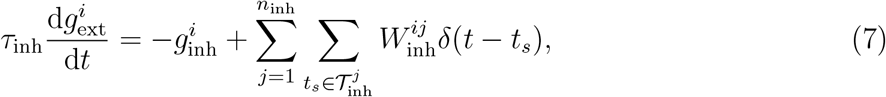

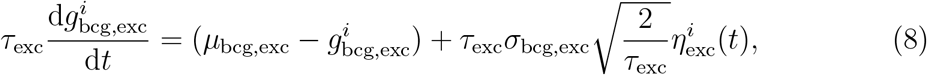

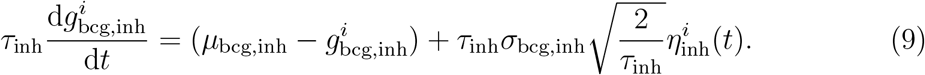

*I*_rec_ is the synaptic current arising from the recurrent connections (exc. to exc., exc. to inh., inh. to exc., inh. to inh.). *I*_ext_ is the excitatory current from external neurons. *I*_bcg_ is the current from synapses from neighboring cortex areas. 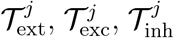 represent the spike times of the *j*-th external, excitatory, and inhibitory neuron respectively. The matrices **W**_ext_, **W**_exc_, **W**_inh_ contain the synaptic connection strengths, 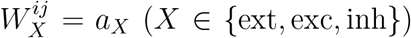 if the *j*-th neuron connects to the *i*-th neuron and 0 otherwise. The input from neighboring cortical areas is modeled as the Ornstein-Uhlenbeck process with means *μ*_bcg,exc_ and *μ*_bcg,inh_ and standard deviations of the limiting distributions *σ*_bcg,exc_ and *σ*_bcg,inh_ [20, 21]. We set the values of the background activity to match the moments of an exponential Poisson shot noise with rates *λ*_bcg,exc_ = 0.5 kHz and *λ*_bcg,inh_ = 0.125 kHz [22]:

**Table 1:**
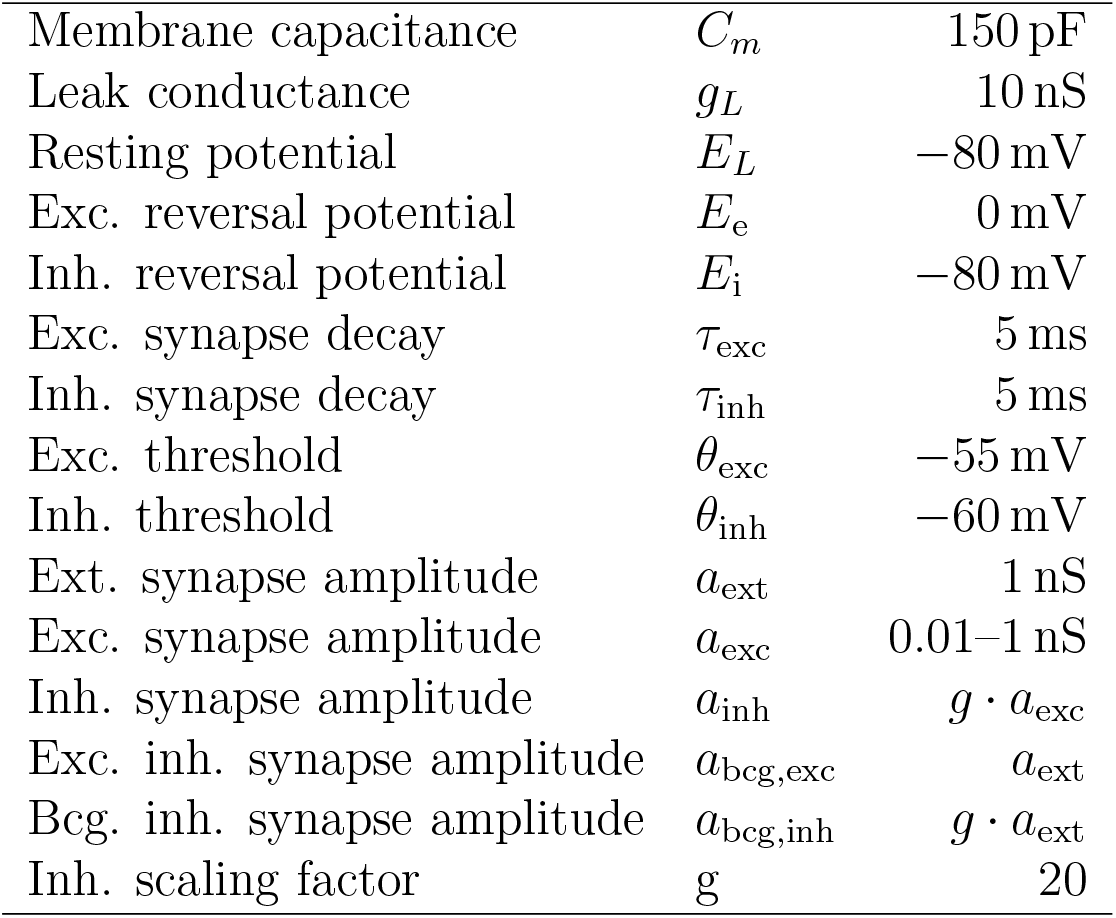
Parameters of the LIF model

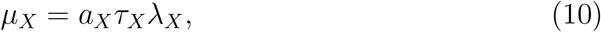

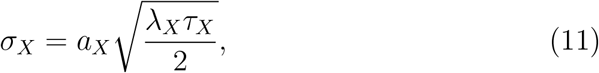

where *X* represents the excitatory or inhibitory background activity.

When the membrane potential *V* crosses the firing threshold (*θ*_exc_, *θ*_inh_) a spike is fired and the membrane potential is reset to *E*_*L*_.

The network consisted of *n*_ext_ = 1000 neurons in the external population, *n*_exc_ = 800 neurons in the excitatory population, and *n*_inh_ = 200 neurons in the inhibitory population. The connections were set randomly with connection probability for the recurrent connections (exc. to exc., exc. to inh., inh. to inh., inh. to exc.) set to *P*_rec_ = 20% and the connection probability from the external population (ext. to exc. and ext. to inh., *P*_ext_) was varied from to 1% to 100% (Fig. 1A). We created the connection matrices **W**_*X*_ by generating a matrix of random uniformly distributed numbers **R**_*X*_ from the interval [0, 1) and set 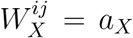 if 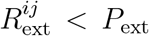 or 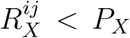 for *X* ∈ {exc, inh}. The random matrix **R**_ext_ was the same for all values of *P*_ext_. In simulations where we controlled for the effects caused by a random number of connections from the external population, we fixed the number of connections by setting only the *k* = *n*_ext_*P*_ext_ elements in each row of *W*_ext_ non-zero, in the location of the *k* largest elements of the *i*-th row of **R**_ext_.

**Figure 1:**
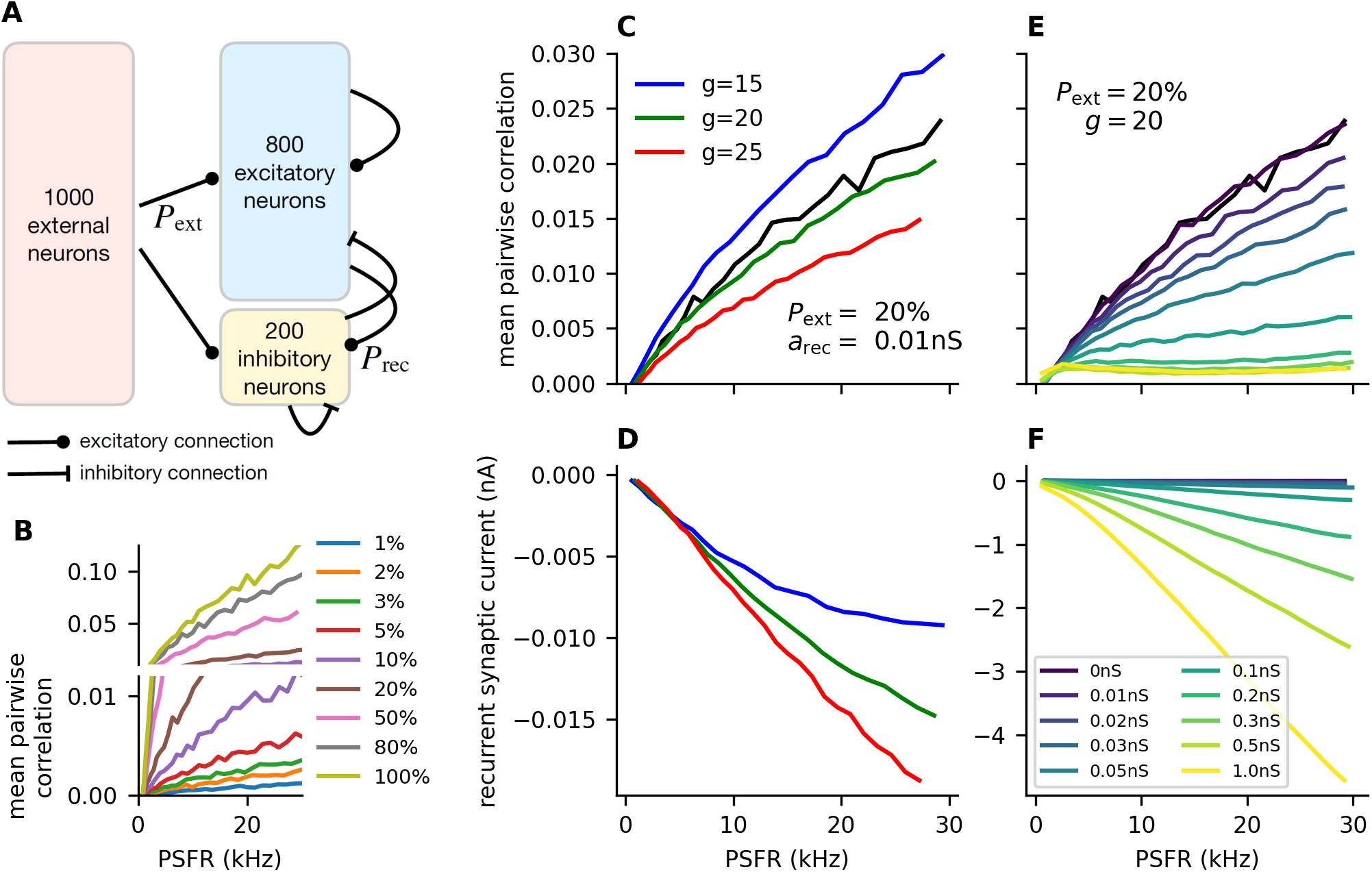
Inhibitory feedback decreases noise correlations. **A:** Schematic illustration of the simulated neural network. Poisson neurons in the external population make random connections to neurons in the excitatory and inhibitory subpopulations. The connection probability *P*_ext_ ∈ [0.01, 1] is varied to achieve different levels of shared external input to the neurons. The neurons in the inhibitory (inh.) and excitatory (exc.) subpopulations make recurrent connections (exc. to exc., exc. to inh., inh. to inh., inh. to exc.) with probability *P*_rec_ = 20%. The strength of those connections is parametrized by *a*_rec_ (Tab. 1). **B:** Mean pairwise correlations between any two neurons in the exc. and inh. subpopulations plotted against the mean output of the network for different values of *P*_ext_ in a feedforward network (*a*_rec_ = 0 nS). **C:** Mean pairwise correlations as in **B**, for different values of *g* (ratio of inhibitory-to-excitatory synaptic strength), *a*_rec_ = 0.01 nS. **D:** Total current from recurrent synapses for different values of *g*, as in *C*. **E-F:** Same as in **C-D**, but with fixed *g* = 20 and different values of *a*_rec_.

The simulations were carried out using the Brian 2 package [23] in Python with a 0.1 ms time step.

### 2.2. Obtaining the input-output relationship of the network

We considered the total number of action potentials *n* from the excitatory and inhibitory subpopulations in time window Δ*T* = 1 s as the output of the network. We modeled the stimulus as the input from the thalamic neurons, parametrized by the mean input rate to a single neuron:

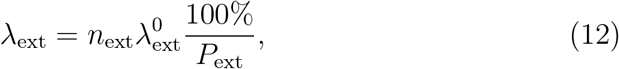

where 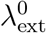 is the firing rate of a single neuron in the external population and 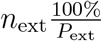 is the mean number of pre-synaptic external neuron for each neuron in the excitatory and inhibitory populations. For each set of parameters (*a*_rec_ and *P*_ext_ pair) we determined the input 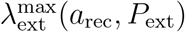 for which the output reached 30 kHz. In order to obtain the input-output relationship, we discretized the input space into 30 equidistant stimulus intensities: 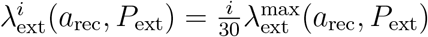, where *i* = 0, …, 30. With a fixed network connectivity, we simulated the network 1080 times for each 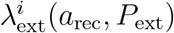.

We then fitted 7th-degree polynomial functions to the mean output of the network as a function of the stimulus *λ*_ext_ and to the Fano factor as a function of the mean output, where Fano factor is defined as:

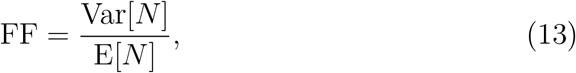

where *N* is a random variable representing the number of output action potentials *n*. The weights of the polynomial fit were set as 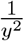, where *y* is the independent variable. We then discretized the input space to 1000 equidistant stimulus intensities and estimated the mean output *μ* and Fano factor FF for each intensity from the polynomial functions. We then estimated the input-output relationship, defined by the conditional probability distribution *f* (*n* |*λ*_ext_) as a lognormal distribution for each *λ*_ext_, with corresponding parameters to match the estimated mean and Fano factor:

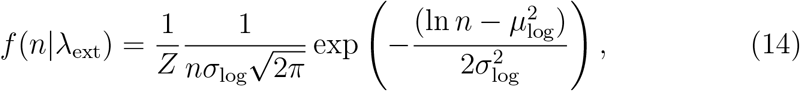

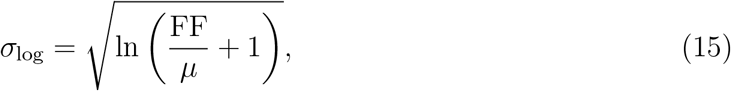

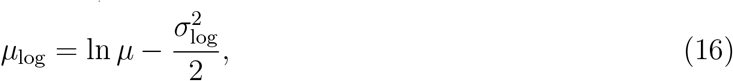

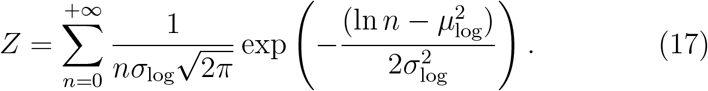

In this way, we avoided the sampling bias when calculating information measures from the data [24].

### 2.3. Metabolic cost of neural activity

In our calculations, we focus on the energy in the form of ATP molecules required to pump out Na^+^ ions. We take into account the Na^+^ influx due to excitatory post-synaptic currents, Na^+^ influx during action potentials, and Na^+^ influx to maintain the resting potential. To this end, we follow the calculations in [2] and [3], which we modify for our neuronal model.

We assume the standard membrane capacitance per area as *c*_*m*_ = 1 μF*/*cm^2^ and the cell diameter as *D* = 69 μm, giving the total capacitance *C*_*m*_ = *πD*^2^*c*_*m*_ = 150 pF. Therefore, to depolarize a neuron by Δ*V* = 100 mV the minimum charge influx is Δ*VC*_*m*_ = 1.5 × 10^−11^ C and the minimum number of Na^+^ ions 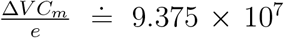, where *e ≐* 1.6 × 10^−19^ C is the elementary charge. The minimal number of Na^+^ ions is then quadrupled to get a more realistic estimate of the Na^+^ influx due to the simultaneous opening of the K^+^ channels [2]. The Na^+^ influx must be then pumped out by the Na^+^*/*K^+^-ATPase, which requires one ATP molecule per 3 Na^+^ ions. The cost of a single action potential can be then estimated as 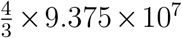 ATP = 1.25 × 10^8^ ATP. However, about 80% of the metabolic costs associated with an action potential are expected to come from the propagation of the action potential through the neuron’s axons. Therefore, we estimate the total cost as 6.25× 10^8^ ATP.

Next, we assume that the excitatory synaptic current is mediated by the opening of Na^+^ and K^+^ channels with reversal potentials *E*_Na_ = 90 mV and *E*_K_ = −105 mV. For the excitatory synaptic current, the following must hold

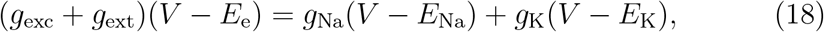

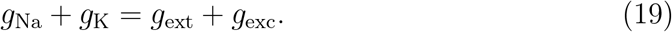

Therefore:

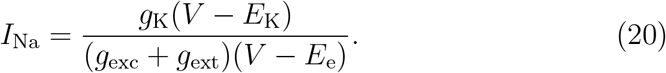

The sodium entering with the sodium current *I*_Na_ must be pumped out by the Na^+^*/*K^+^-ATPase, and therefore we calculate the cost of the synaptic current as 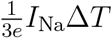 ATP, where Δ*T* is the time interval over which we are measuring the cost.

Each input to the network (parametrized by *λ*_ext_) is then associated with a cost, which we express as

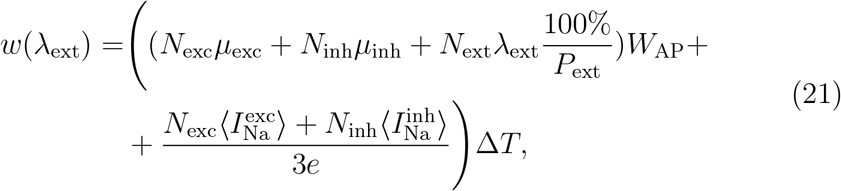

where *μ*_exc_ = *μ*_exc_(*λ*_ext_), *μ*_inh_ = *μ*_inh_(*λ*_ext_) are the mean firing rates of a single excitatory and inhibitory neuron (given the input *λ*_ext_), 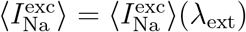 and 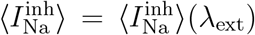 are the average excitatory synaptic currents in a single excitatory and inhibitory neuron.

### 2.4. Measuring the information content

We treat the neural network as a memoryless information channel [25, 26]. The firing rates of the external population *λ*_ext_ are the input to the channel, and the number of action potentials *n* that the excitatory population fires in the time window Δ*T* = 1 s is the output of the channel. The input is then described by a random variable Λ and the output by a random variable *N*. The mutual information between the input and the output *I*(Λ; *N*) is calculated as

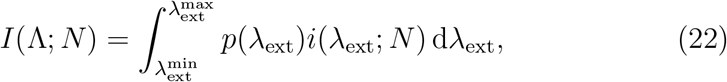

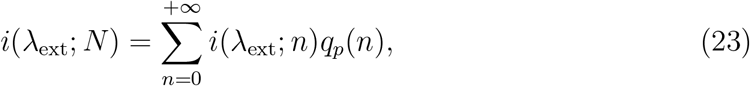

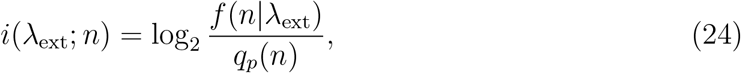

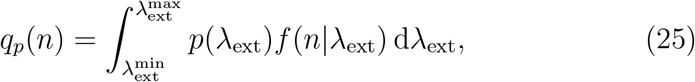

where *f* (*n*|*λ*_ext_) is the probability distribution of *N* given that Λ = *λ*_ext_, *p*(*λ*_ext_) is the input probability distribution, *i*(*λ*_ext_; *n*) is the amount of information that an observation of *n* spikes gives us about the stimulus *λ*_ext_, *i*(*λ*_ext_; *N*) is then the average amount of information we get from the input *λ*_ext_, *q*_*p*_(*n*) is the marginal output probability distribution.

Given the input probability distribution *p*(*λ*_ext_), we can calculate the average metabolic cost as

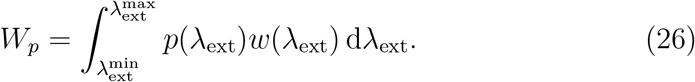

The capacity-cost function *C*(*W*) is then the lowest upper bound on the amount of mutual information (in bits) achievable given the constraint that *W*_*p*_ *< W* :

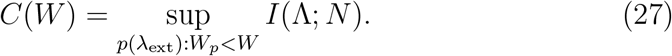

The information-metabolic efficiency *E* is then the maximal amount of mutual information per molecule of ATP between the input and the output:

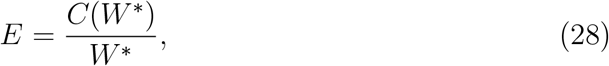

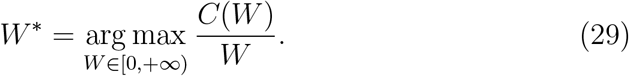

The capacity-cost function can be obtained numerically with the Blahut-Arimoto algorithm [27], or the information-metabolic efficiency can be conveniently obtained directly with the Jimbo-Kunisawa algorithm [28, 29].

#### 2.4.1. Low noise approximation of constrained information capacity

If the trial-to-trial variability is very low, a lower bound on the capacity-cost function can be found [30, 31]. We used this low-noise approximation to gain analytical insight into the importance of different properties of the neural system for information-metabolically efficient information transmission. In the low noise approximation, the optimal input distribution maximizing the mutual information constrained by metabolic expenses *W* is given by

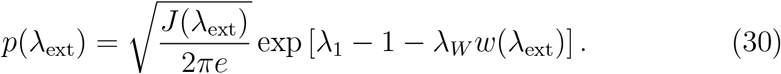

where *J* (*λ*_ext_) is the Fisher information and *λ*_1_ and *λ*_*W*_ are the Lagrange multipliers which can be obtained from the normalization condition:

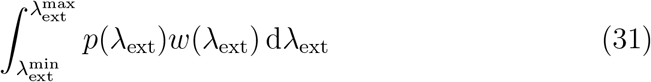

and the average metabolic cost constraint (Eq. 26). In the Gaussian approximation, the Fisher information is given by

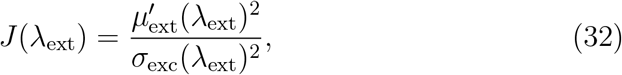

where *σ*_exc_(*λ*_ext_) is the standard deviation of the spike counts at input intensity *λ*_ext_. The low noise estimate on the capacity-cost function is then

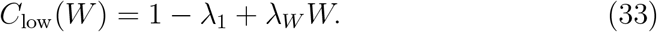

## 3. Results

### 3.1. Constrained information maximization in a simple linear model

In order to gain an insight into what affects the information-metabolic efficiency of a neural population, we first solve the problem for a simple linear system. The mean response of the system is given by *γ*(*λ*_ext_) = *gλ*_ext_, where *λ*_ext_ is the stimulus and *g* is the gain of the system. The Fano factor (Eq. 13) is constant, and we assume that the output is continuous and normally distributed. Therefore, the input-output relationship is described by

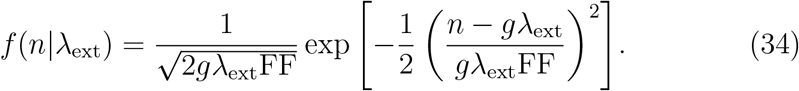

and the Fisher information (Eq. 32) is

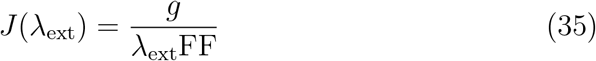

Next, we assume that the cost of the activity *w*(*λ*_ext_) depends linearly on the input:

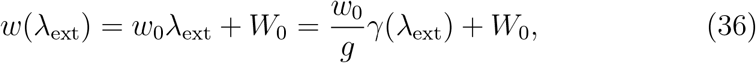

where *W*_0_ is the cost of the resting state.

The probability distribution derived from the low-noise approximation (Eq. 30) is then

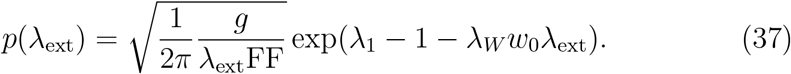

After applying the normalization conditions (Eqs. 26 and 31) and using Eq. (33) we obtain the lower bound on the capacity-cost function:

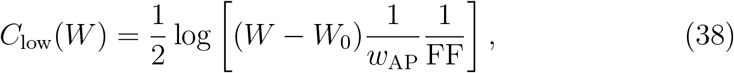

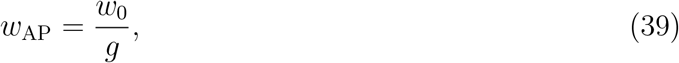

where *w*_AP_ is the cost of increasing the output intensity by one action potential.

The gain *g*, cost scaling *w*_0_, and Fano factor FF cannot be considered constant for real neural populations. However, Eq. (38) provides an insight into the importance of these properties, which we will study numerically for a more realistic neural system.

In the following, we use

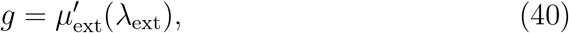

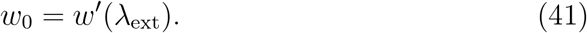

### 3.2. Inhibitory feedback decorrelates the neural activity

With the increasing probability of shared input (*P*_ext_), the mean pairwise correlation between the firing output of neurons increases (feedforward network, Fig. 1B). These correlations could be removed by recurrent connections, as long as the synaptic currents from the recurrent connections were inhibition dominated. We set the excitatory recurrent synaptic amplitude as *a*_exc_ = 0.01 nS to create a small perturbation from the feedforward network and varied the scaling *g* determining the amplitude of inhibitory synapses (*a*_inh_ = *ga*_exc_) from 15 to 25. Correlations between neurons were decreased for *g* ≥20 (Fig. 1C), which was also associated with stronger negative net current from the recurrent synapses (Fig. 1D). For the network considered further in our work, we set *g* = 20. Simultaneously increasing the strength of the recurrent synapses with fixed *g* led to a further decrease of the correlations among the neurons (Fig. 1E) while further decreasing the net current from the recurrent synapses (Fig. 1F).

### 3.3. Trial-to-trial variability of single neurons vs. a population

In an inhibition-dominated network, the input needed from the external population in order to evoke a given average firing rate has to be higher than in the case of the feedforward network. The resulting increase in synaptic noise leads to higher trial-to-trial variability in the LIF model (Fig. 2 A-C; see also [32]).

**Figure 2:**
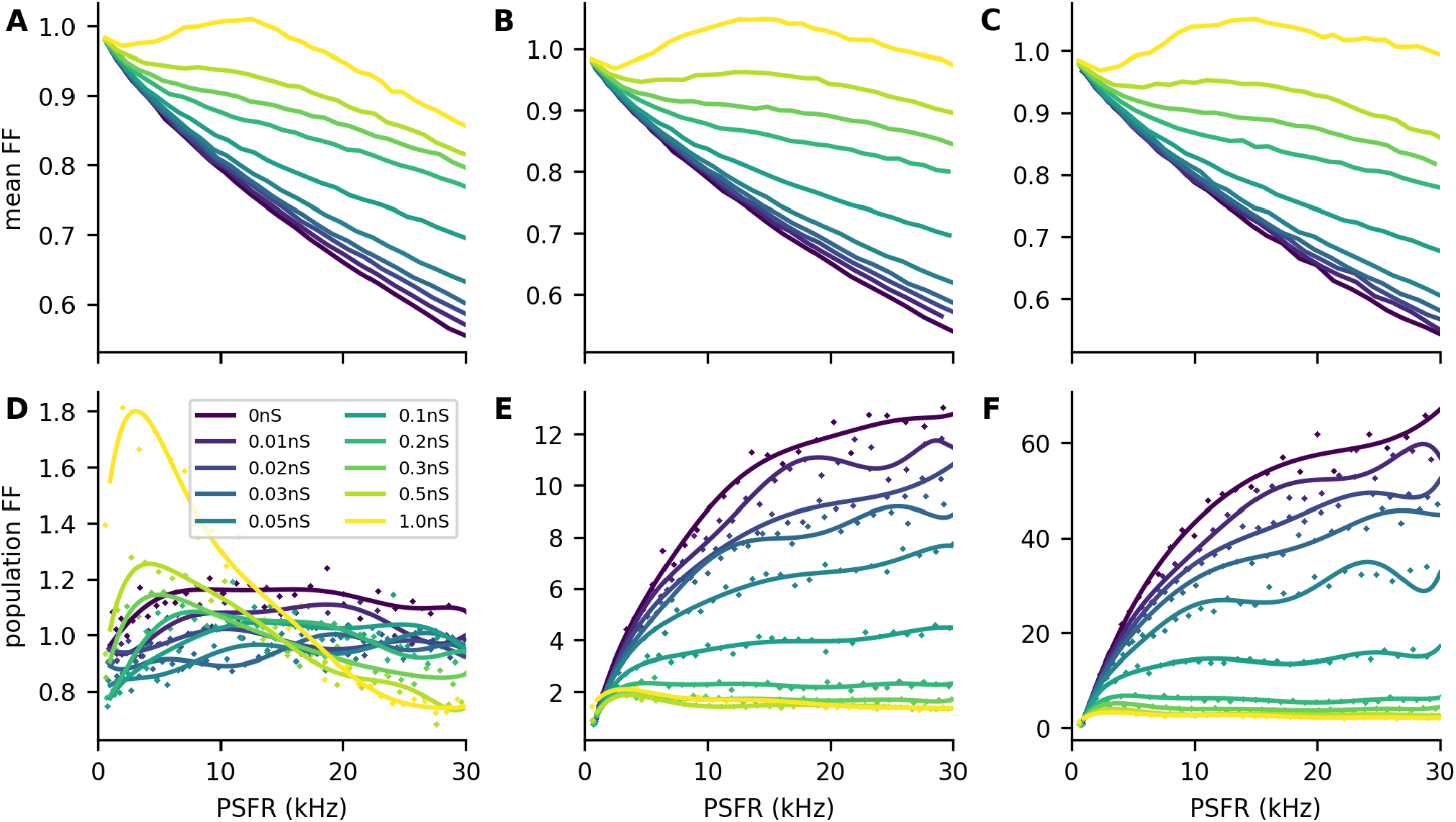
Fano factor of single neurons and of populations. **A-C:** Mean Fano factor of individual neurons for different values of *P*_ext_: 0.01 (**A**), 0.2 (**B**), 1 (**C**). The strength of the recurrent synapses (*a*_rec_) is colorcoded. The mean Fano factor increases with the strength of the recurrent synapses. **D-F:** Same as in **A-C** but for the Fano factor of the population activity. The points represent the population Fano factor obtained from the simulation, and the lines represent the fit with a 7th-degree polynomial. For *P*_ext_ = 0.01, the increase in trial-to-trial variability of individual neurons (**A**) can have a stronger effect on the population Fano factor than decreasing the pairwise correlations, resulting in an increase of the population Fano factor with high values of *a*_rec_ (**D**). For higher values of *P*_ext_, the pairwise correlations greatly increase the population Fano factor, which then decreases with increasing *a*_rec_.

In the case of the total population activity, however, the pairwise correlations between the neurons have a significant effect on the trial-to-trial variability. By denoting the random variable representing the number of spikes of the *i*-th neuron observed during time window Δ*T* as *N*_*i*_, we get for the Fano factor of the population activity:

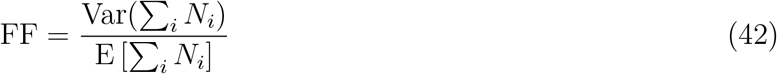

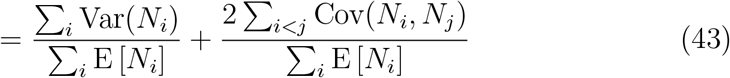

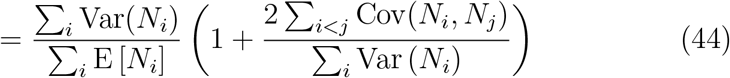

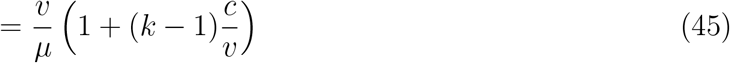

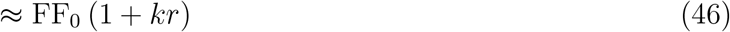

where *c* is the mean pairwise covariance, *v* the mean variance of a neuron, *μ* is the mean number of spikes in Δ*T, k* is the number of neurons, and *r* is the Pearson correlation coefficient. The last approximation holds for neurons with identical variances and pairwise covariances [16]. It provides an insight into how the pairwise correlations and Fano factor of individual neurons affect the Fano factor of the total activity. If the correlations or number of neurons are small (*r·k* ≪ 1), the decorrelation by strengthening the recurrent synapses does not significantly decrease the population Fano factor. Instead, the population Fano factor may increase due to the increase of the Fano factor of individual neurons (Fig. 2D, *P*_ext_ = 1%). If greater correlations are induced due to the shared input to the network, the correlations have a dominating effect on the population Fano factor, which can then be greatly decreased by strengthening the recurrent synapses and in turn decreasing the pairwise correlations (Fig. 2E-F).

### 3.4. Inhibitory feedback is metabolically costly

#### 3.4.1. Stronger recurrence strength increases the cost of the resting state

The cost of the resting state is an important factor for informationmetabolic efficiency [10]. In our network, increasing the recurrence strength decreased the spontaneous activity of the neurons, due to inhibition dominating the recurrent currents. However, the simultaneous increase in the strength of the recurrent excitatory synapses increased the cost of the excitatory synaptic currents (Fig. 3A-C).

**Figure 3:**
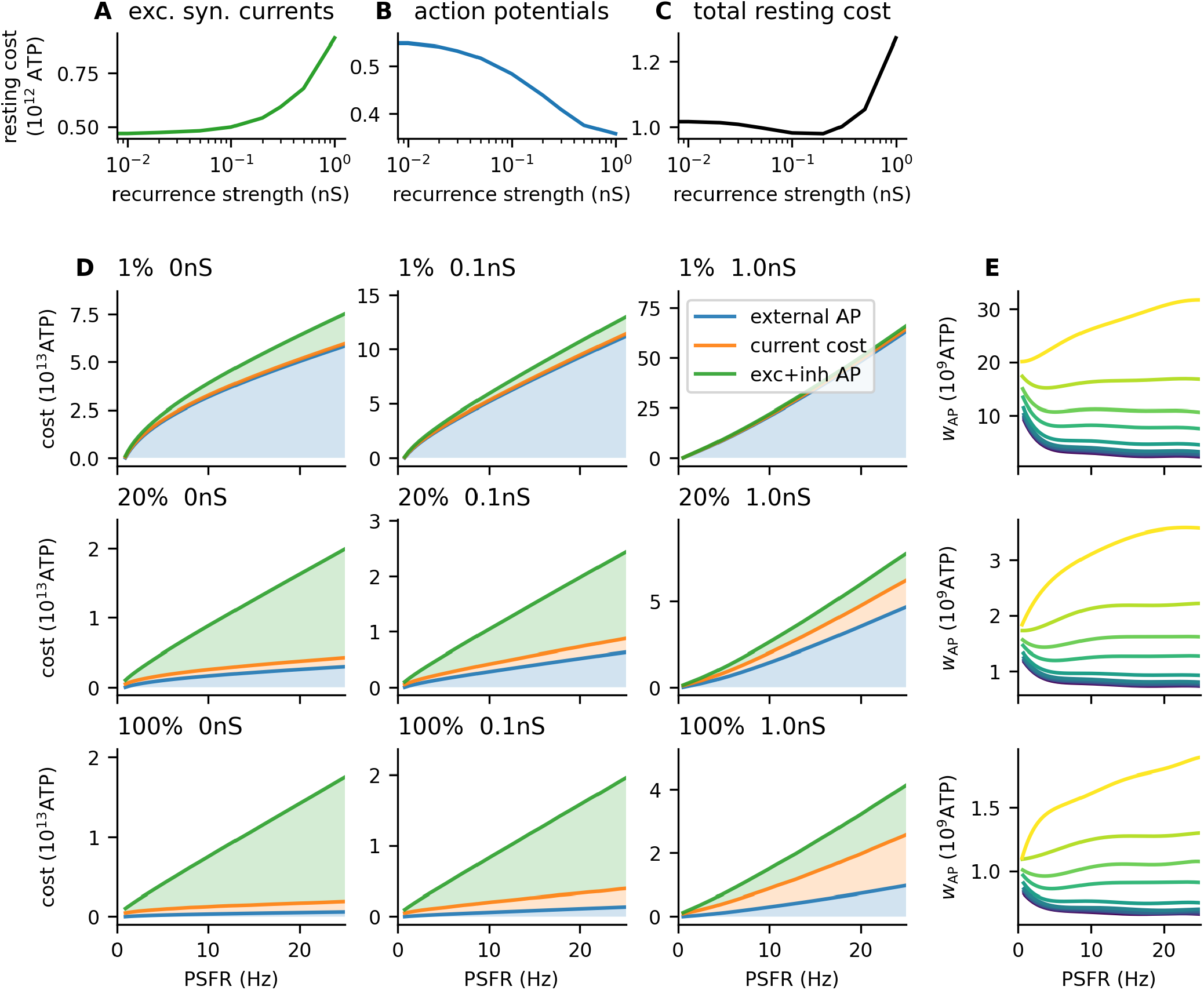
Metabolic cost of the network activity. **A-C:** Cost at resting state (*λ*_ext_ = 0). **A:** Cost of the excitatory synaptic currents from the background activity and excitatory action potentials evoked by the background activity. **B:** Cost of the action potentials (both excitatory and inhibitory) evoked by the background activity. **C:** Total resting cost obtained by summing **A** and **B. D:** The total cost of the network activity is plotted against the output of the network (the total post-synaptic firing rate). Filled areas represent individual contributions of each cost component: cost of action potentials from the external population, cost of the excitatory synaptic currents, and cost of the post-synaptic (evoked) action potentials. As *P*_ext_ increases, the contribution of external action potentials to the overall cost decreases. With increasing *a*_rec_, the contribution of excitatory synaptic currents increases. **E:** The cost of increasing the mean input by one action potential (*w*_AP_, Eq. 39) is significantly lower for higher *P*_ext_. However, although the difference between *P*_ext_ = 1% and *P*_ext_ = 20% is approximately 10-fold, the difference between *P*_ext_ = 20% and *P*_ext_ = 100% is only approximately 2-fold, as the cost of the external population starts to contribute less to the overall cost.

#### 3.4.2. Inhibitory feedback decreases gain

Because the net current from recurrent synapses is hyperpolarizing, with stronger recurrent synapses, a stronger excitatory current is necessary to bring the neuron to a given post-synaptic firing rate, and higher pre-synaptic firing rates are necessary. Therefore, the gain *g* of the network decreases, and with increasing *a*_rec_ the cost of synaptic currents and the cost of external activity increase (Fig. 3D-E).

### 3.5. Shared input decreases gain

The number of synapses from the external population for each neuron in the excitatory and inhibitory subpopulations follows a binomial distribution:

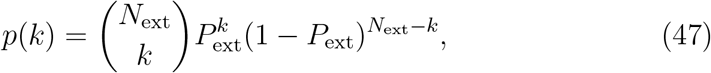

with the mean number of synapses given by *N*_ext_ · *P*_ext_ and variance *N*_ext_ · *P*_ext_(1 − *P*_ext_). We scaled the firing rate of the individual neurons in the external population as 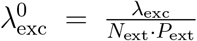. Therefore the mean output to a single neuron was always *λ*_ext_, independently of *P*_ext_ and the variance of the input across neurons was 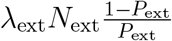.

Given the convexity of the single neuron tuning curve in the analyzed input range (Fig. S1) that out of two inputs with an identical mean *λ*_ext_, but different variances across neurons, the input with the higher variance will lead to a higher average firing rate. Assuming that the input across neurons follows a normal distribution with mean *λ*_ext_ and variance *σ*^2^ and that the single neuron tuning curve can be approximated by an exponential function in the form of *c*_1_ exp(*c*_2_*x*), where *x* is the input intensity to the single neuron, we obtain the mean firing rate:

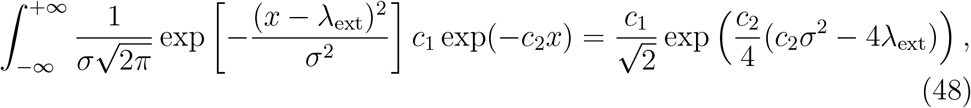

which grows with the standard deviation of the input.

Accordingly, we observed that networks with higher *P*_ext_ needed higher *λ*_ext_ in order to produce the same mean PSFR as networks with lower *P*_ext_ (Fig. 4A-C). Moreover, the mean Fano factor of individual neurons increased with increasing *P*_ext_ (Fig. 4D-F). This effect could be mostly removed by fixing the number of connections from the external population to each neuron in the excitatory and inhibitory populations to *P*_ext_*N*_ext_ (Fig. S2).

**Figure 4:**
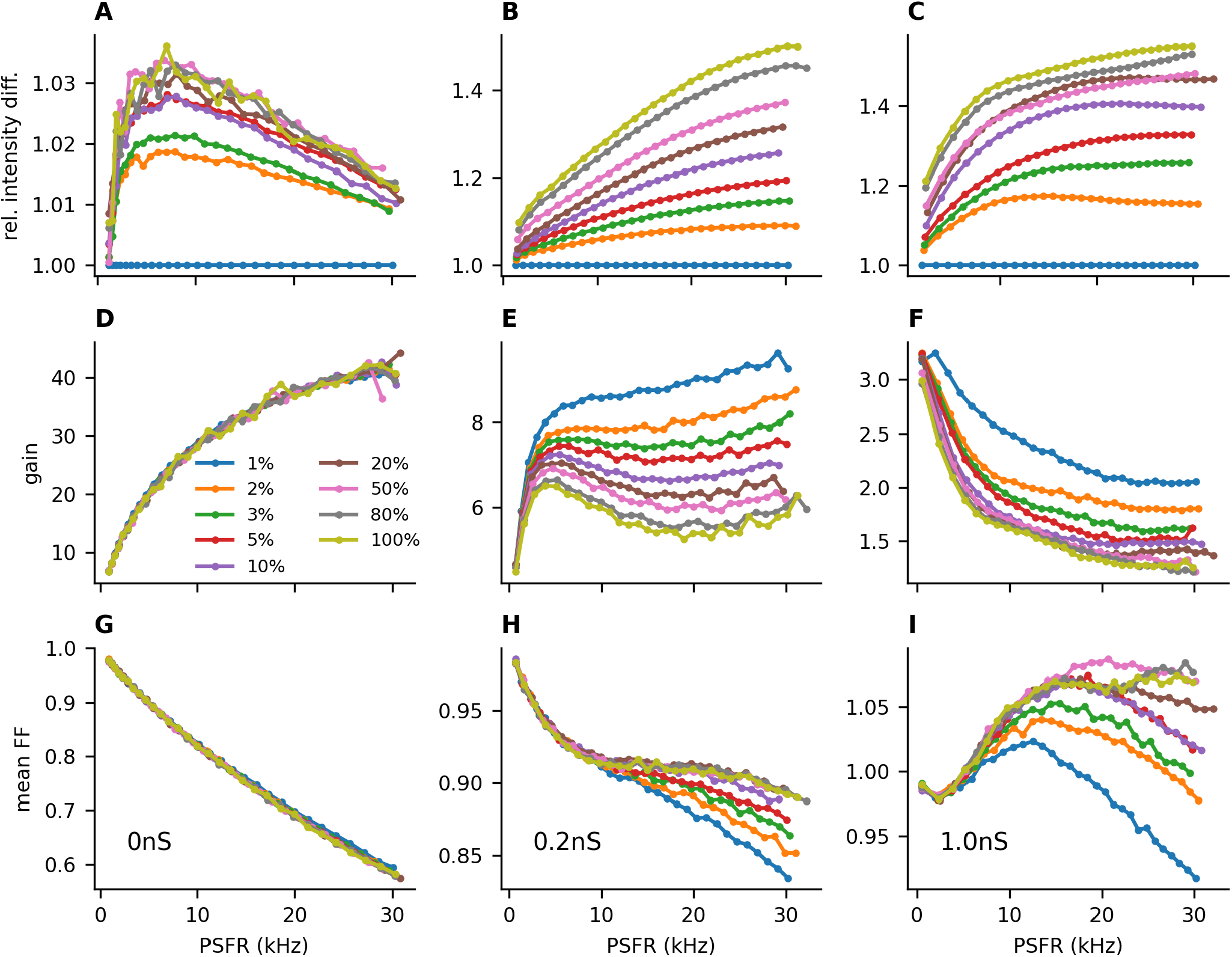
Shared input decreases the gain and increases the individual trial-to-trial variability. **A-C:** The input intensity *λ*_ext_ needed to evoke a given firing rate (*x*-axis) with different connection probabilities *P*_ext_ relative to the input intensity for *P*_ext_ = 1%. **A:** *a*_rec_ = 0 nS, **B:** *a*_rec_ = 0.2 nS, **C:** *a*_rec_ = 1 nS. For higher *P*_ext_, higher values of *λ*_ext_ are needed to achieve the same post-synaptic firing rates as with lower values of *P*_ext_. This effect becomes more pronounced in stronger recurrent synapses (**E-F**). **D-F:** Gain of the network (Eq. 40). A higher *P*_ext_ leads to a lower gain of the population activity. **G-I:** Higher values of *P*_ext_ also increase the Fano factor of individual neurons.

### 3.6. Optimal regimes for metabolically efficient information transmission

We illustrated that the recurrence strength 1) increases the metabolic cost of the neural activity and 2) decreases the trial-to-trial variability of the population response by decreasing the correlations between the neurons. Similarly, the increased probability of a synapse from an external neuron to a neuron in the excitatory or inhibitory population decreases the cost of the neural activity but increases the trial-to-trial variability of the population response by increasing the noise correlations. To find the balance between the cost of the network activity (Eq. 26) and the mutual information between the input and the output (Eq. 22), we calculated the information-metabolic efficiency, which maximizes the ratio of the mutual information to the cost of the network activity (Eq. 28).

For low values of *P*_ext_ (≤10%), increasing the strength of the recurrent input did not lead to an increase in the information-metabolic efficiency. For higher values of *P*_ext_ the information-metabolic efficiency was maximized for *a*_rec_ between 0.1 nS and 0.5 nS (Fig. 5A-B), meaning that the strength of the recurrent excitatory synapses was 2× to 5× lower that the strength of the synapses from the external population.

**Figure 5:**
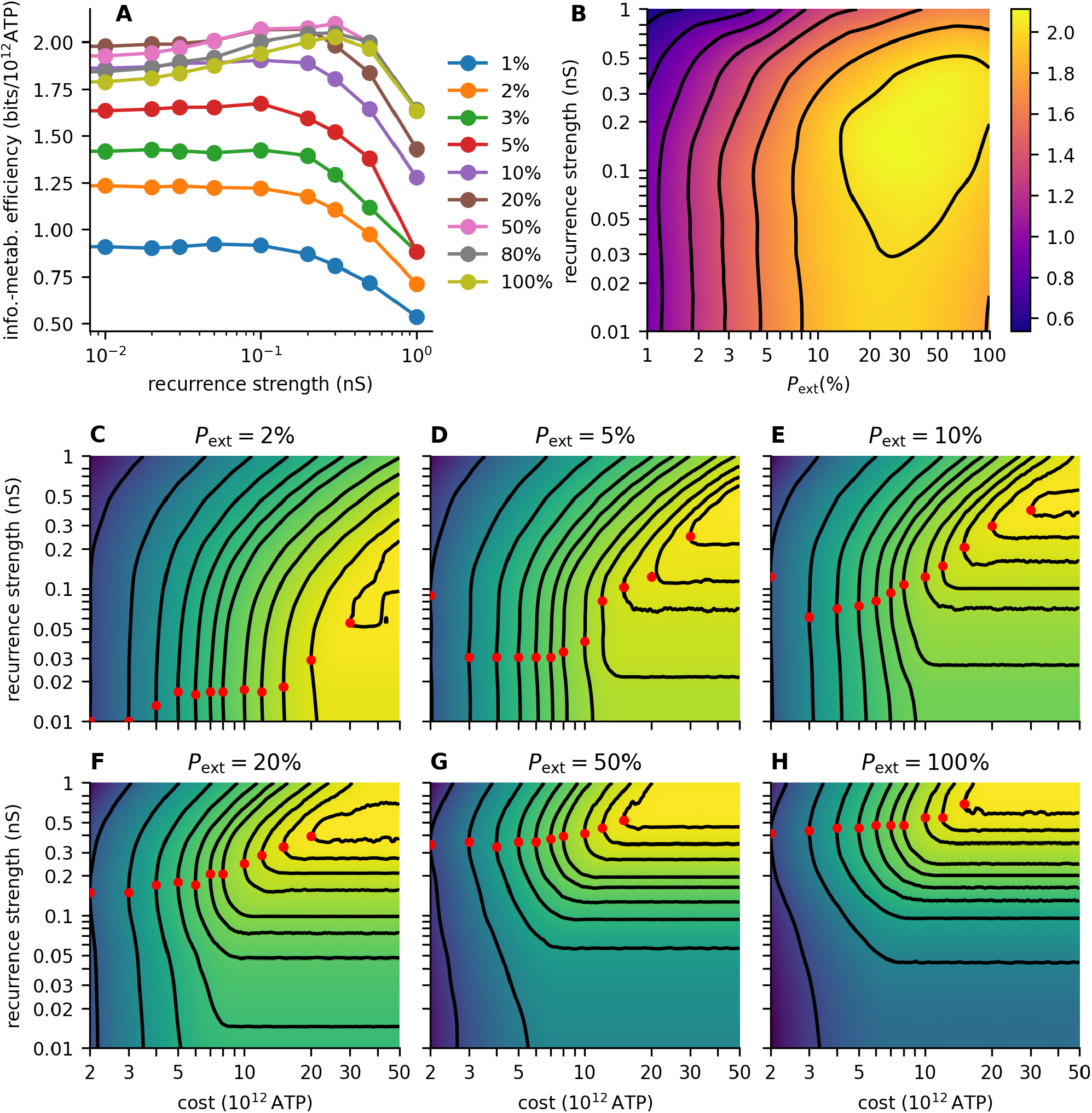
Information transmission with cost constraints. **A:** Information-metabolic efficiency *E* (Eq. 28) for different values of recurrence strength *a*_rec_. *P*_ext_ is color-coded. **B:** Contour plot of the informationmetabolic efficiency. Contours are at 0.6, 0.8, 1, 1.2, 1.4, 1.6, 1.8, and 2 bits/s. **C-H:** Contour plots showing the capacity-cost function *C*(*W*) (Eq. 27) with dependence on the recurrence strength *a*_rec_ for different values of *P*_ext_. The contours show the maximal capacities constraint at different values of *W* (see Tab. 2 for the costs and capacity values at the contours). The heatmaps in **B-H** were calculated using piece-wise cubic 2D interpolation (SciPy interpolator CloughTocher2DInterpolator [33]) from the grid calculated with *P*_ext_ values 1%, 2%, 3%, 5%, 10%, 20%, 50%, 80%, 100% and *a*_rec_ values 0, 0.01, 0.02, 0.03, 0.05, 0.1, 0.2, 0.3, 0.5, and 1 nS.

Moreover, varying *P*_ext_ had a significant effect on the information-metabolic efficiency across all values of *a*_rec_. Namely, low values of *P*_ext_ resulted in lower values of information-metabolic efficiency across all values of *a*_rec_, showing that shared input from the external population is beneficial for metabolically efficient information transmission. Overall, the highest values of information-metabolic efficiency (*E* ≥ 2 bit*/*10^12^ATP) were reached for *a*_rec_ between 0.05 nS and 0.5 nS and *P*_ext_ between 20% and 100% (Fig. 5B).

We analyzed the effect of the resting cost (Fig. 3A-C) by setting the resting cost in all cases equal to *W*_0_, the resting cost of the feedforward network. This did not have a significant effect on the information-metabolic efficiencies (Fig. S3).

Neural circuits might not necessarily maximize the ratio of information to cost. Instead, neurons and neural circuits could modulate their properties to maximize information transmission with the available energy resources [5]. For example, neurons in the mouse visual cortex have been shown to decrease the conductance of their synaptic channels after food restriction [34].

Accordingly, we studied how the optimal strength of recurrent synapses changes with the available resources. We calculated the optimal value of *a*_rec_ for different values of available resources (3, 4, 5, 6, 7, 8, 10, 12, 15, 20, 30, and 40 ×10^12^ ATP). In Fig. 5C-H, we plotted *C*(*W* ; *a*_rec_), the capacity-cost function (Eq. 27) extended by one dimension with *a*_rec_. For each cost *W*, the optimal *a*_rec_ is highlighted, and the corresponding contour of *C*(*W*) is shown (see Tab. 2 for the values of *C*(*W*)). With decreasing *W*, the optimal value of *a*_rec_ typically decreases. This effect is more robust with high values of *P*_ext_, because the contours are more curved at the optimum.

We calculated the extended capacity-cost functions using input distributions obtained from the low-noise approximation. To verify that the low noise approximation applies in the case of the studied system, we compared these results to the information-metabolic efficiency obtained with the Jimbo-Kunisawa algorithm. The relative difference did not exceed 10% and did not have a significant impact on the information-metabolic efficiency heatmap structure (Fig. S4).

**Table 2:**
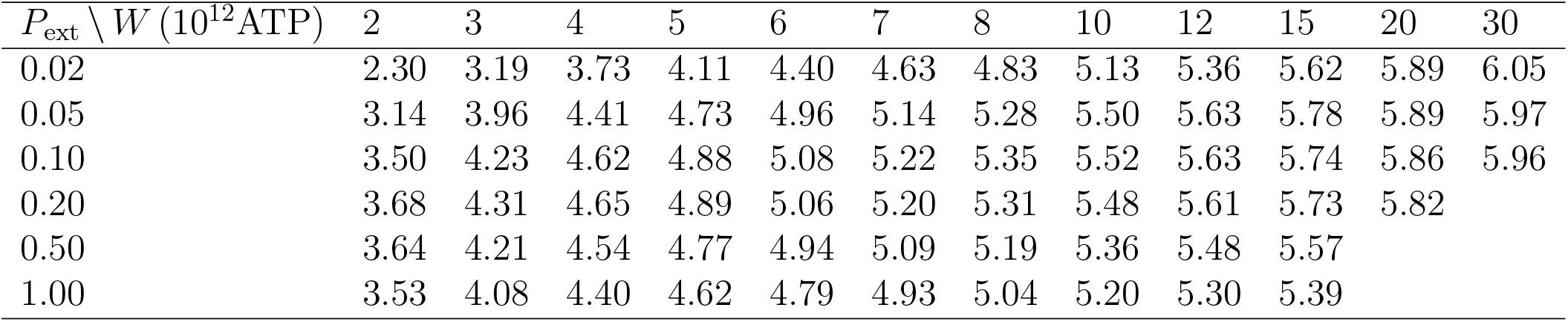
capacity-cost function values (in bits)

## 4. Discussion

Populations in the cortex that transmit information by their summed (or averaged) activity can be considered as a low noise information channel, due to the decrease in trial-to-trial variability [30]. The decrease in the trial-to-trial variability of the response will be lower in the presence of positive noise correlations [16]. Positive noise correlations can be reduced by inhibitory feedback, which, however, increases the cost of the neural activity [10].

In our work, we studied the balance between increasing the transmitted information by decreasing the noise correlations and the associated increase in the cost of the activity. We showed that in a linear system, if the Fano factor of the population activity and the ratio 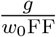 (*g* is the gain of the system, or slope of the stimulus-response curve, *w*_0_ is the slope of the stimulus-cost curve) remain constant, the cost-constrained capacity will remain constant as well.

We proceeded to calculate the stimulus-response relationship and the metabolic cost for a more biologically realistic neural system. In the studied system, the population Fano factor could not be considered constant. Instead, correlations between neurons increased with the mean output of the system, and the mean Fano factor of single neurons was also dependent on the mean output of the system, leading to complex dependence of the population Fano factor on the mean output of the system (Fig. 2D-F). We showed that reducing the noise correlations by strengthening the inhibitory feedback decreases the population Fano factor for large values of *P*_ext_ but may increase the population Fano factor for low values of *P*_ext_ due to an increase of the mean single neuron Fano factor FF_0_.

We illustrated the effect of inhibition-dominated recurrence and shared input on the metabolic cost of neural activity. An increased strength of recurrence increased the cost of excitatory synaptic currents due to the stronger excitatory synapses and stronger input from the external population, as well as the cost of the activity of the external population. A higher connection probability from the external population (higher shared input probability) led to a decrease in the external population activity cost, as the overall activity of the external population could be lower to result in the same mean input to the post-synaptic neurons. On the other hand, due to less variable input to single neurons with high values of *P*_ext_, a higher mean input was required across all neurons to evoke the same mean post-synaptic activity.

We found that high values of *P*_ext_ were beneficial for metabolically efficient information transmission, despite the increased noise correlations. For high values of *P*_ext_, increasing the recurrence strength was also beneficial, suggesting that the two mechanisms - decreasing input cost by high connection probability from the external population *P*_ext_ and decreasing noise correlations by recurrent activity may act together to produce a metabolically efficient code.

Increasing the recurrence strength can lead to a 10% to 15% increase in the information-metabolic efficiency. The magnitude of the increase is dependent on the cost of the action potentials. If the cost of synaptic currents is negligible compared to the cost of the action potentials, there would be a higher benefit in increasing the inhibitory feedback since the increases in the cost of the synaptic current could also be neglected.

Although neurons in the cortex also connect to neighboring areas of the cortex and not only within the studied subpopulation, we did not consider the cost of synaptic currents evoked in neurons not involved in our simulation. We assume that such synaptic currents would be part of the background activity of a different area. Therefore, if we included these costs and considered multiple cortical areas, we would have included the background activity cost multiple times.

In our model of the cortical area, we considered two neural subpopulations: excitatory and inhibitory. Each subpopulation was homogeneous, but we set the threshold of the inhibitory neurons lower to mimic the behavior of fast-spiking inhibitory neurons. The difference between excitatory, regular spiking neurons and inhibitory, fast-spiking neurons is often described not only by differences in the threshold but also in differences in the adaptation properties [35, 36, 37]. In our case, we did not consider adaptation for simplicity because estimating the information capacity of a neural system with adaptation is computationally considerably more difficult [10].

In our work, we assumed that the neural circuit maximizes the mutual information between the input and the output neurons while minimizing the cost of the neural activity. Such an approach does not provide any information about how the information is encoded. It only calculates the limit on the amount of information that can be reliably transmitted. Yet, the principles of mutual information maximization have proven very useful in explaining the properties of neural systems. For example, the tuning curves of blowfly’s contrast-sensitive neurons are adapted to the distribution of contrasts encountered in the natural environment [38]; the power spectrum of distribution of odor in pheromone plumes follows the power spectrum predicted for an optimal input to olfactory receptor neurons [39]; distributions of post-synaptic firing rates of single neurons during *in-vivo* recordings follow distributions predicted from cost-constrained mutual information maximization [40, 41, 42].

By assuming a particular coding scheme, it is possible to place further constraints on the complexity of information encoding, with the assumption that complex codes are not an efficient way to transmit information [43, 44]. We did not attempt this in our study. However, it would be interesting to study whether inhibitory feedback decreases or increases the encoding complexity.

We have shown that a cortical area can adapt to the amount of available energy resources. When resources are scarce, information transmission can be adapted by weakening the synaptic weights, thus expending fewer resources to reduce the noise correlations. Such a mechanism is implemented in the mouse visual cortex [34]. **(author?)** showed that in food-restricted mice, the orientation tuning curves of individual orientation-sensitive neurons in the visual cortex become broader due to weakened synaptic conductances.

In our work, we studied the properties of a neuronal population instead of single neurons. In particular, we considered a population encoding the stimulus intensity instead of the stimulus identity, such as orientation. An extension this model to a situation in which stimulus identity is encoded and shared input is introduced due to the overlap of receptive fields would be interesting.

Neurons recorded *in-vivo* typically exhibit a Fano factor close to 1.0 and constant over a broad range of post-synaptic firing rates [45, 46, 18]. In the optimal regimes with stronger recurrent synapses, the Fano factor decreased only very slowly over the studied range of post-synaptic firing rates (up to 30 Hz in a single neuron). With weaker synaptic strengths, the Fano factor of a single neuron decreases rapidly with an increasing post-synaptic firing rate. Our model predicts that fewer available resources would lead to weaker recurrent synapses. This hypothesis is straightforward to test by calculating the Fano factors during stimulus presentation (both population and single neuron) in food-restricted animals and comparing them to controls. We expect that the population Fano factor will increase (alternatively, the noise correlations will increase) with food scarcity, and single neuron Fano factors will decrease.

## Acknowledgments

This work was supported by Charles University, project GA UK No. 1042120. TB benefited from a fellowship from the Plant Health and Environment Division of INRAE.

---

**Figure S1:**
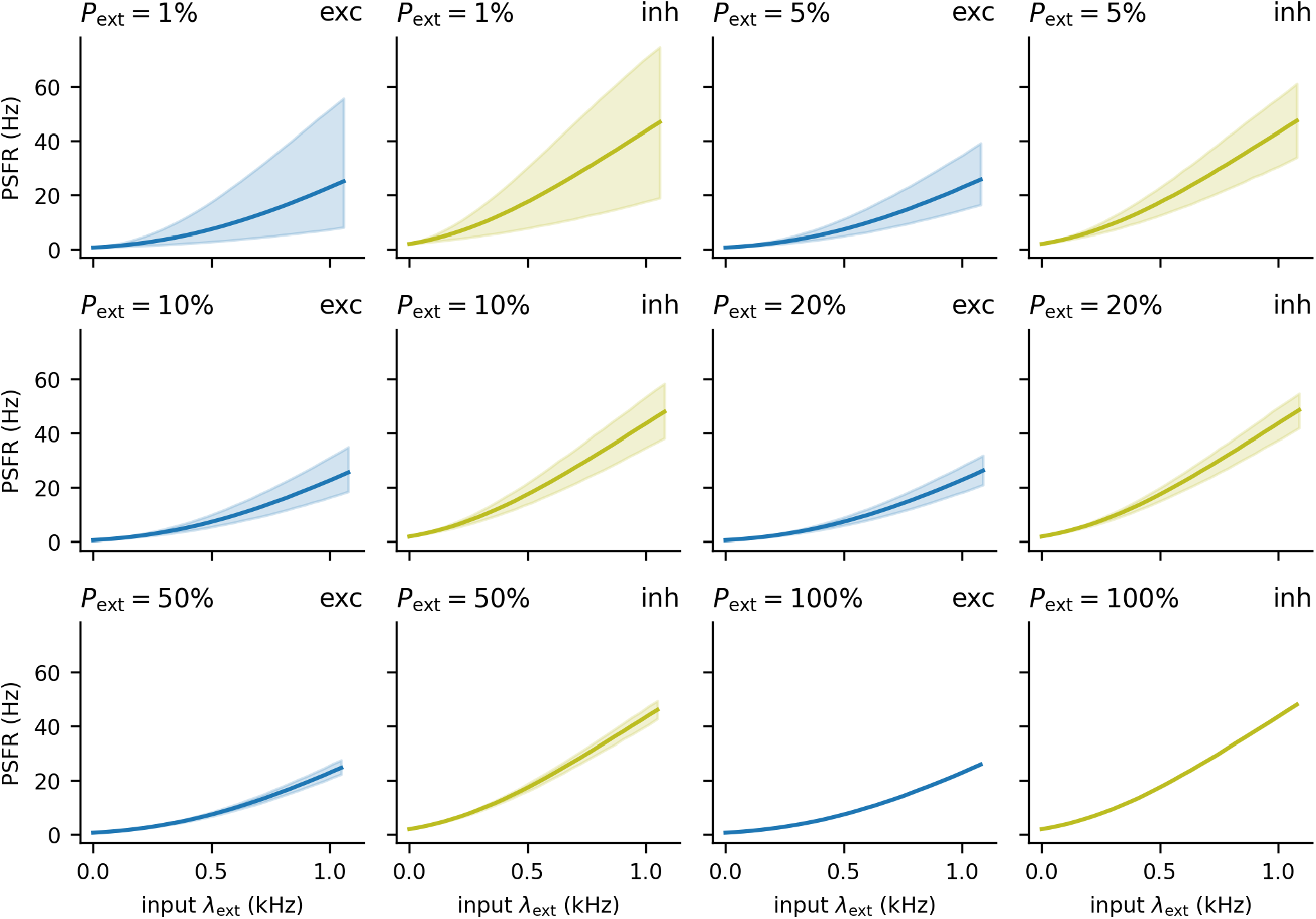
Input-output relationship of single neurons. To exclude the network effects, we plotted the tuning curves for the feedforward network separately for the excitatory (blue) and inhibitory (yellow) neurons. The thick line represents the median response across the neurons, which shows that their tuning curves are convex in the studied range. The shaded area shows the spread of the tuning curves across neurons (2.5 to 97.5 percentile). With low values of *P*_ext_, the tuning curves across neurons vary significantly and are skewed to the higher firing rates.

**Figure S2:**
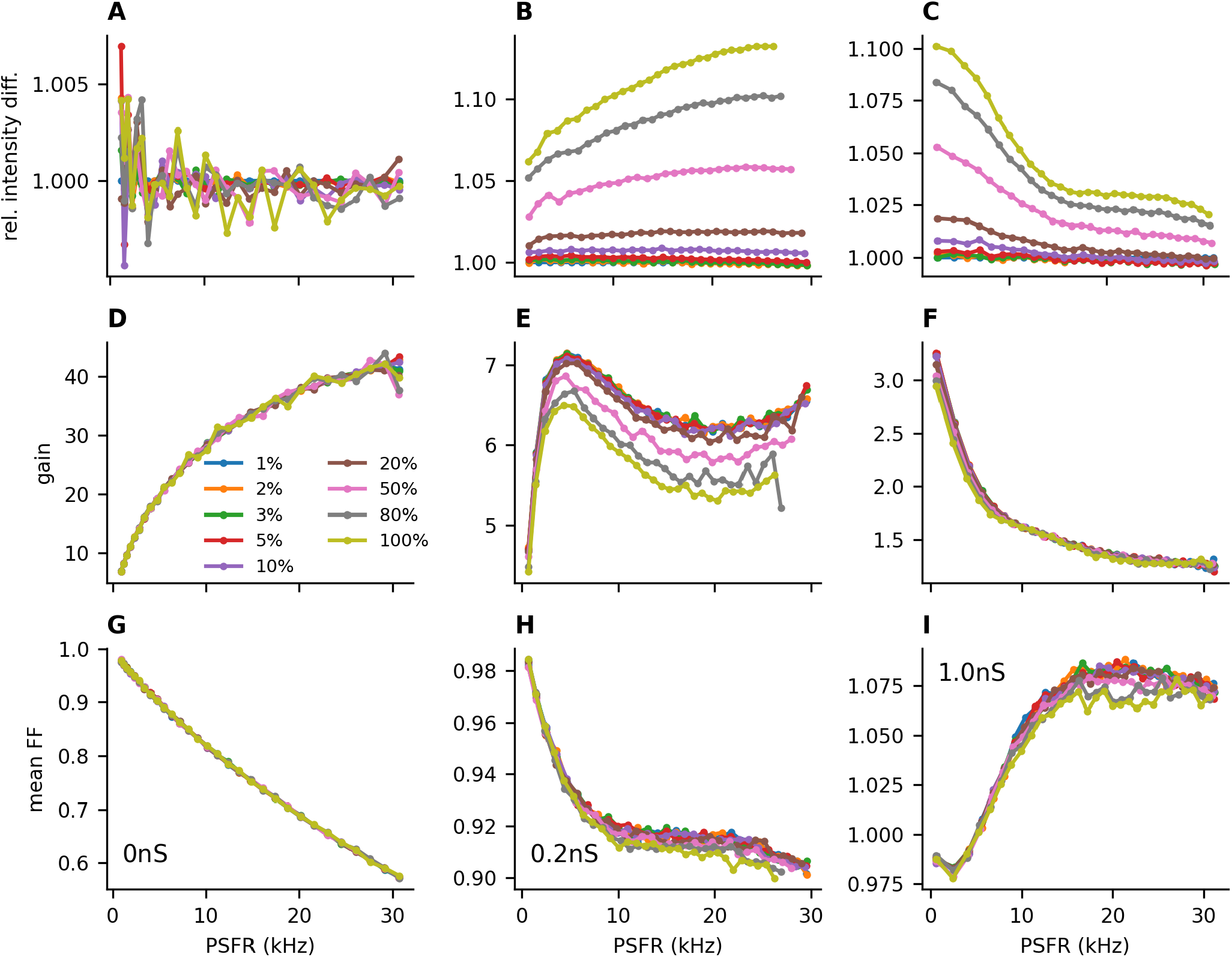
Fixing the number of external connections to each neuron. Same as Fig. 4, but exactly 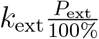 external neurons connected to each excitatory and inhibitory neuron. This removed a large part of the dependence on *P*_ext_ seen in Fig. 4.

**Figure S3:**
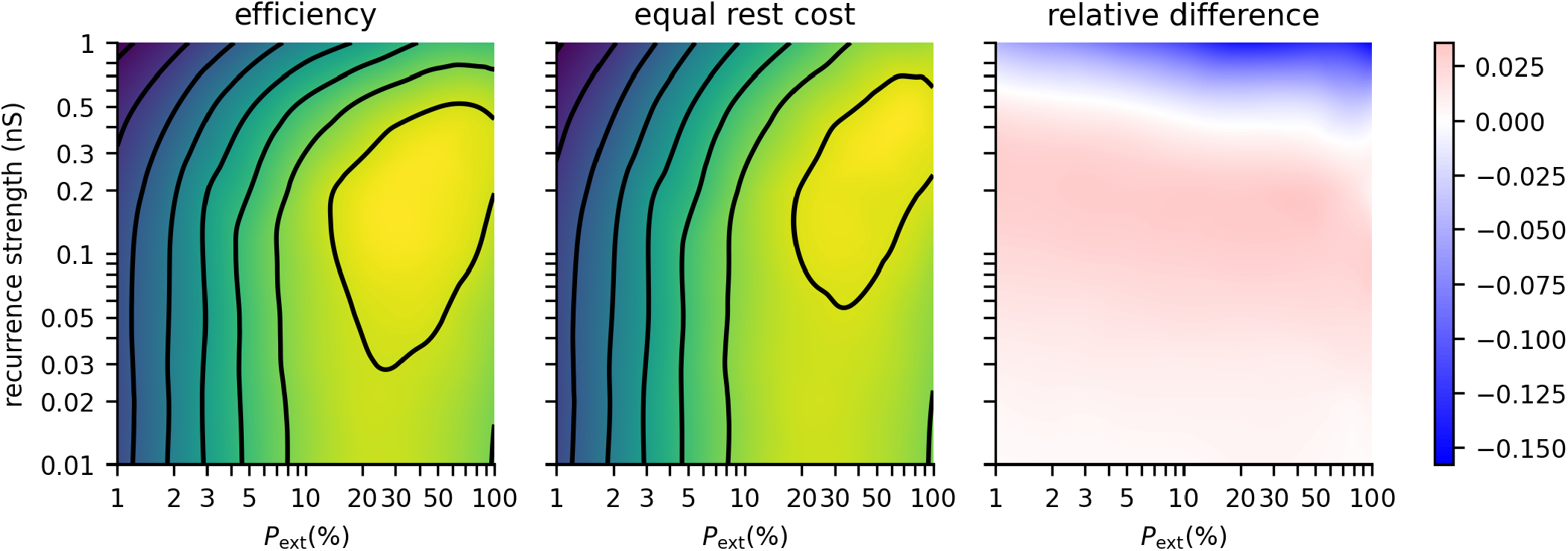
Effect of equalizing the resting cost on the information-metabolic efficiency. We observed that the cost of the resting state was different for different recurrence strengths *a*_rec_ (Fig. 3A-C). This could potentially explain the higher information-metabolic efficiency *E* (Eq. 28) for intermediate values of *a*_rec_ and its decrease for high values of *a*_rec_. To quantify the effect of the resting cost, we set the resting cost in each case to the resting cost of the feedforward network *W*_0_(*a*_rec_ = 0). The differences in the cost of the resting state did not have a qualitative effect on the conclusions. **A:** The same contour plot as in Fig. 5B. **B:** Contour plot with equalized resting costs (contours as in Fig. 5B: 0.6, 0.8, 1, 1.2, 1.4, 1.6, 1.8 and 2 bits/s). **C:** Heatmap of the relative differences.

**Figure S4:**
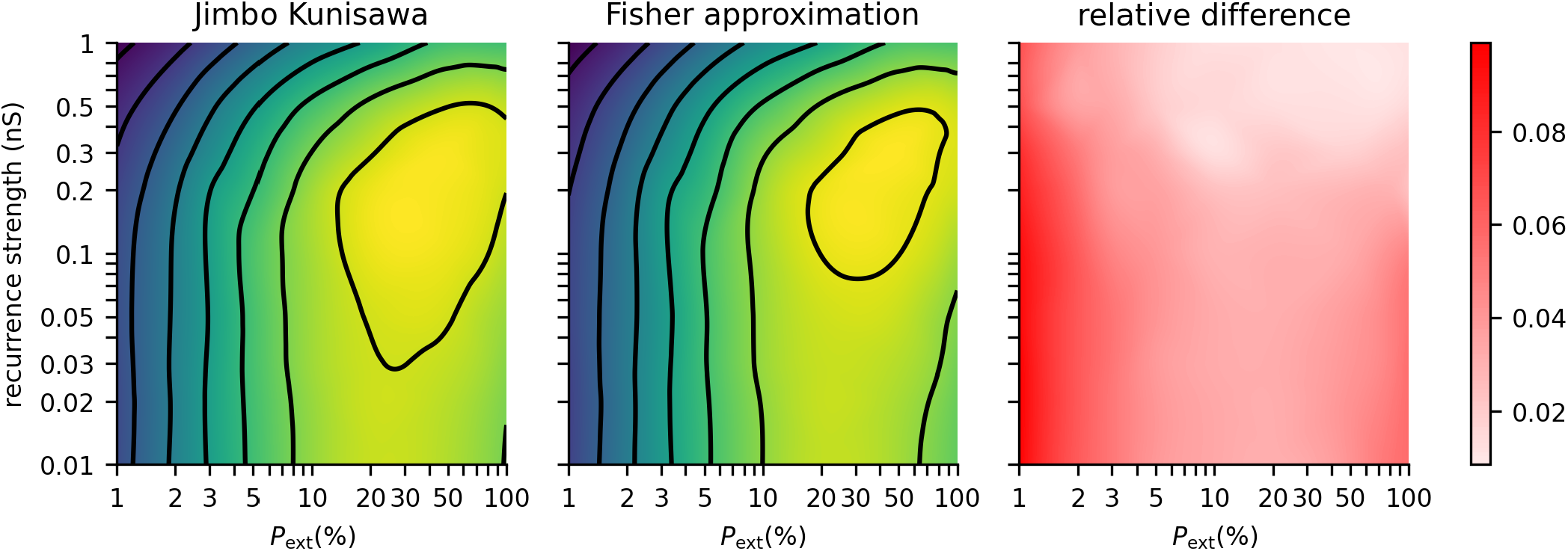
Accuracy of information-metabolic efficiency approximation. To calculate the capacity-cost functions, we calculated the mutual information using Eq. (22) with the input probability distribution calculated from Eqs. (30) and (32). Here we compare the information-metabolic efficiencies calculated with the approximation and the Jimbo-Kunisawa algorithm. **A:** The same contour plot as in Fig. 5B with information-metabolic efficiencies calculated with the Jimbo-Kunisawa algorithm. **B:** Information-metabolic efficiencies calculated with the Fisher-information-based input distribution. **C:** Heatmap of the relative differences. Note that the approximation can only reach values lower than the actual information-metabolic efficiency.

## Notes

### Competing Interest Statement

The authors have declared no competing interest.

### Summary of Updates

Formatting change, proofreading, and language editing.

## References

[1] H. B. Barlow, Possible Principles Underlying the Transformations of Sensory Messages, in: Sensory Communication, The MIT Press, 1961, pp. 217–234. doi:10.7551/mitpress/9780262518420.003.0013.

[2] D. Attwell, S. B. Laughlin, An Energy Budget for Signaling in the Grey Matter of the Brain, J. Cereb. Blood Flow Metab. 21 (10) (2001) 1133–1145. doi:10.1097/00004647-200110000-00001.

[3] J. J. Harris, R. Jolivet, D. Attwell, Synaptic energy use and supply, Neuron 75 (5) (2012) 762–777.

[4] W. B. Levy, R. A. Baxter, Energy Efficient Neural Codes, Neural Comput. 8 (3) (1996) 531–543. doi:10.1162/neco.1996.8.3.531.

[5] V. Balasubramanian, D. Kimber, M. J. Berry II, Metabolically Efficient Information Processing, Neural Comput. 13 (4) (2001) 799–815. doi:10.1162/089976601300014358.

[6] S. Laughlin, Energy as a constraint on the coding and processing of sensory information, Curr. Opin. Neurobiol 11 (4) (2001) 475–480. doi:10.1016/S0959-4388(00)00237-3.

[7] J. E. Niven, S. B. Laughlin, Energy limitation as a selective pressure on the evolution of sensory systems, J. Exp. Biol. 211 (11) (2008) 1792–1804. doi:10.1242/jeb.017574.

[8] L. Yu, Y. Yu, Energy-efficient neural information processing in individual neurons and neuronal networks, J. Neurosci. Res. 95 (11) (2017) 2253–2266. doi:10.1002/jnr.24131.

[9] B. Sengupta, S. B. Laughlin, J. E. Niven, Balanced Excitatory and Inhibitory Synaptic Currents Promote Efficient Coding and Metabolic Efficiency, PLoS Comput. Biol. 9 (10) (2013) e1003263. doi:10.1371/journal.pcbi.1003263.

[10] T. Barta, L. Kostal, The effect of inhibition on rate code efficiency indicators, PLoS Comput Biol. 15 (12) (2019) e1007545. doi:10.1371/journal.pcbi.1007545.

[11] C. Monier, F. Chavane, P. Baudot, L. J. Graham, Y. Frégnac, Orientation and Direction Selectivity of Synaptic Inputs in Visual Cortical Neurons, Neuron 37 (4) (2003) 663–680. doi:10.1016/s0896-6273(03)00064-3.

[12] N. Brunel, Dynamics of Sparsely Connected Networks of Excitatory and Inhibitory Spiking Neurons, J. Comput. Neurosci. 8 (2000) 183–208. doi:https://doi.org/10.1023/A:1008925309027.

[13] A. Renart, J. de la Rocha, P. Bartho, L. Hollender, N. Parga, A. Reyes, K. D. Harris, The Asynchronous State in Cortical Circuits, Science 327 (5965) (2010) 587–590. doi:10.1126/science.1179850.

[14] T. Tetzlaff, M. Helias, G. T. Einevoll, M. Diesmann, Decorrelation of Neural-Network Activity by Inhibitory Feedback, PLoS Comp. Biol. 8 (8) (2012) e1002596. doi:10.1371/journal.pcbi.1002596.

[15] A. Bernacchia, X.-J. Wang, Decorrelation by Recurrent Inhibition in Heterogeneous Neural Circuits, Neural Comput. 25 (7) (2013) 1732–1767. doi:10.1162/NECO_a_00451.

[16] L. F. Abbott, P. Dayan, The Effect of Correlated Variability on the Accuracy of a Population Code, Neural Comput. 11 (1) (1999) 91–101. doi:10.1162/089976699300016827.

[17] B. B. Averbeck, P. E. Latham, A. Pouget, Neural correlations, population coding and computation, Nat. Rev. Neurosci. 7 (5) (2006) 358–366. doi:10.1038/nrn1888.

[18] M. N. Shadlen, W. T. Newsome, The Variable Discharge of Cortical Neurons: Implications for Connectivity, Computation, and Information Coding, J. Neurosci. 18 (10) (1998) 3870–3896. doi:10.1523/jneurosci.18-10-03870.1998.

[19] R. Moreno-Bote, J. Beck, I. Kanitscheider, X. Pitkow, P. Latham, Pouget, Information-limiting correlations, Nature Neuroscience 17 (10) (2014) 1410–1417. doi:10.1038/nn.3807.

[20] G. E. Uhlenbeck, L. S. Ornstein, On the Theory of the Brownian Motion, Phys. Rev. 36 (5) (1930) 823–841. doi:10.1103/physrev.36.823.

[21] A. Destexhe, M. Rudolph, J. M. Fellous, T. J. Sejnowski, Fluctuating synaptic conductances recreate in vivo-like activity in neocortical neurons., Neuroscience 107 (1) (2001) 13–24.

[22] K. Rajdl, P. Lansky, Stein’s neuronal model with pooled renewal input, Biol. Cybern. 109 (3) (2015) 389–399. doi:10.1007/s00422-015-0650-x.

[23] M. Stimberg, R. Brette, D. F. Goodman, Brian 2, an intuitive and efficient neural simulator, eLife 8 (8 2019). doi:10.7554/eLife.47314.

[24] A. Treves, S. Panzeri, The Upward Bias in Measures of Information Derived from Limited Data Samples, Neural Comput. 7 (2) (1995) 399–407. doi:10.1162/neco.1995.7.2.399.

[25] C. Shannon, A mathematical theory of communication, Bell system technical journal 27 (1948).

[26] J. A. T. Thomas M. Cover, Elements of Information Theory, 2nd Edition, Wiley Series in Telecommunications and Signal Processing, Wiley-Interscience, 2006.

[27] R. Blahut, Computation of channel capacity and rate-distortion functions, IEEE Trans Inf Theory 18 (4) (1972) 460–473. doi:10.1109/tit.1972.1054855.

[28] M. Jimbo, K. Kunisawa, An iteration method for calculating the relative capacity, Information and Control 43 (2) (1979) 216–223. doi:10.1016/s0019-9958(79)90719-8.

[29] P. Suksompong, T. Berger, Capacity Analysis for Integrate-and-Fire Neurons With Descending Action Potential Thresholds, IEEE Trans Inf Theory 56 (2) (2010) 838–851. doi:10.1109/tit.2009.2037042.

[30] L. Kostal, P. Lansky, Information capacity and its approximations under metabolic cost in a simple homogeneous population of neurons, Biosystems 112 (3) (2013) 265–275. doi:10.1016/j.biosystems.2013.03.019.

[31] L. Kostal, P. Lansky, M. D. McDonnell, Metabolic cost of neuronal information in an empirical stimulus-response model, Biol. Cybern. 107 (3) (2013) 355–365. doi:10.1007/s00422-013-0554-6.

[32] T. Barta, L. Kostal, Regular spiking in high-conductance states: The essential role of inhibition, Phys Rev E 103 (2) (2021) 022408. doi:10.1103/PhysRevE.103.022408.

[33] P. Virtanen, R. Gommers, T. E. Oliphant, M. Haberland, T. Reddy, D. Cournapeau, E. Burovski, P. Peterson, W. Weckesser, J. Bright, S. J. van der Walt, M. Brett, J. Wilson, K. J. Millman, N. Mayorov, A. R. J. Nelson, E. Jones, R. Kern, E. Larson, C. J. Carey, SciPy 1.0: fundamental algorithms for scientific computing in Python, Nature Methods 17 (3) (2020) 261–272. doi:10.1038/s41592-019-0686-2.

[34] Z. Padamsey, D. Katsanevaki, N. Dupuy, N. L. Rochefort, Neocortex saves energy by reducing coding precision during food scarcity, Neuron 110 (2) (2022) 280–296. doi:10.1016/j.neuron.2021.10.024.

[35] R. Kobayashi, Y. Tsubo, S. Shinomoto, Made-to-order spiking neuron model equipped with a multi-timescale adaptive threshold, Front. Comput. Neurosci. 3 (2009) 9. doi:10.3389/neuro.10.009.2009.

[36] Y. Zerlaut, S. Chemla, F. Chavane, A. Destexhe, Modeling mesoscopic cortical dynamics using a mean-field model of conductance-based networks of adaptive exponential integrate-and-fire neurons, J. Comput. Neurosci 44 (1) (2017) 45–61. doi:10.1007/s10827-017-0668-2.

[37] D. Bernardi, G. Doron, M. Brecht, B. Lindner, A network model of the barrel cortex combined with a differentiator detector reproduces features of the behavioral response to single-neuron stimulation, PLOS Comput Biol 17 (2) (2 2021). doi:10.1371/journal.pcbi.1007831.

[38] S. Laughlin, A simple coding procedure enhances a neuron’s information capacity., Z. Naturforsch. [C] 36 (9-10) (1981) 910–912.

[39] L. Kostal, P. Lansky, J.-P. Rospars, Efficient olfactory coding in the pheromone receptor neuron of a moth, PLoS Comput. Biol. 4 (2008) e1000053.

[40] A. Treves, S. Panzeri, E. T. Rolls, M. Booth, E. A. Wakeman, Firing rate distributions and efficiency of information transmission of inferior temporal cortex neurons to natural visual stimuli., Neural Comput. 11 (3) (1999) 601–632.

[41] G. G. de Polavieja, Errors Drive the Evolution of Biological Signalling to Costly Codes, J. Theor. Biol. 214 (4) (2002) 657–664. doi:10.1006/jtbi.2001.2498.

[42] G. G. de Polavieja, Reliable biological communication with realistic constraints, Phys Rev E 70 (6) (12 2004). doi:10.1103/physreve.70.061910.

[43] L. Kostal, R. Kobayashi, Optimal decoding and information transmission in Hodgkin-Huxley neurons under metabolic cost constraints, Biosystems 136 (2015) 3–10. doi:10.1016/j.biosystems.2015.06.008.

[44] L. Kostal, R. Kobayashi, Critical size of neural population for reliable information transmission, Phys. Rev. E (Rapid Commun.) 100 (1) (2019) 050401(R).

[45] M. Gur, A. Beylin, D. M. Snodderly, Response Variability of Neurons in Primary Visual Cortex (V1) of Alert Monkeys, J. Neurosci 17 (8) (1997) 2914–2920. doi:10.1523/JNEUROSCI.17-08-02914.1997.

[46] W. S. Geisler, D. G. Albrecht, Visual cortex neurons in monkeys and cats: Detection, discrimination, and identification, Vis Neurosci 14 (5) (1997) 897–919. doi:10.1017/S0952523800011627.

